# Higher levels of *grp78* and lower levels of *tlr4* anticipate a positive evolution of sepsis and patient survival

**DOI:** 10.1101/133264

**Authors:** Razvan C. Stan, Camila P. Bonin, Rose Porto, Francisco G. Soriano, Maristela M. de Camargo

## Abstract

In sepsis caused by Gram-negative bacteria, modulation of Toll-like receptor 4 (TLR4) activity by modulators such as glucose-regulated protein 78 kDa (GRP78), is believed to shift the equilibrium between pro- and anti-inflammatory downstream signaling cascade. We measured daily mRNA *tlr4* and *grp78* expression levels in peripheral blood of a cohort of septic patients, upon intensive care admission, and modeled these mRNA values based on a sine damping function. We obtained negative correlations between *tlr4* and *grp78* mRNA in the survivor group. In contrast, such relation is lost in the deceased patients. Loss of homeostasis predicted by our model within the initial 5 days of hospitalization was confirmed by death of those patients up to 28 days later. Measuring the correlation patterns of the expression of these two genes serves as a robust means to gauge sepsis progression, requiring only three points of measurement on the first day of hospitalization.

## Introduction

Sepsis syndrome is a major public health concern that involves both the physiological and pathological reactions of the organism to invading pathogens and their toxins. The Sepsis-related Organ Failure Assessment (SOFA) uses at least two of the following clinical criteria for its diagnosis: respiratory rate ≥ 22 mpm, altered mentation, or systolic blood pressure ≤ 100 mm Hg [1]. As the condition worsens, sepsis leads to at least one organ dysfunction, and further develops into septic shock characterized by hypotension unresponsive to fluid therapy. This latter stage is associated with high morbidity and mortality rates [2].

Toll-family members link innate and adaptive immunity [3]. TLR4 senses lipopolysaccharides (LPS) from Gram-negative bacteria but also endogenous ligands from the host (eg. fibrinogen) [4,5]. The signaling cascade thus initiated, and the ultimate mounting of the immune response, requires the involvement of the unfolded protein response (UPR) to ensure overall quality control of multiple proteins needed to be synthesized, folded, and secreted. Exposure to low doses of LPS inhibits the endoplasmic reticulum (ER)-stress response in macrophages and activates the inositol-requiring enzyme 1 (IRE1)-X-box binding protein 1 (XBP1) pathway that augments the initial production of pro-inflammatory cytokines [6]. However, continued LPS stimulation ultimately induces the apoptosis of immune cells, an important pathological alteration observed during sepsis [7]. For instance, abnormal myocardial and lymphocytic apoptosis are associated with an exacerbation of the UPR and are observed in later sepsis stages. This includes an initial upregulation of the anti-apoptotic glucose-regulated protein 78 kDa (GRP78) [8], followed by increased levels of the pro-apoptotic C/EBP homologous protein (CHOP) [9-11]. Consequently, attenuation of ER stress with pharmacological agents reduces the overall amplitude of inflammation, as do the anti-apoptotic therapies that rescue experimental models from peritoneal and respiratory sepsis [12].

More than 170 sepsis biomarkers have been put forth as their detection implicate, among others, complement and coagulation activation, and inflammation [13]. Whereas the early sepsis stage associated with the pro-inflammatory signaling cascade is generally reversible, patients that progress into septic shock are more likely to succumb, in spite of in-hospital therapy [14]. Consequently, as the response to sepsis varies over time, the period during which any specific biomarker is useful will change accordingly, complicating its reliability for prognosis [13, 15]. The early involvement of surface TLR4 in mediating systemic responses to both invading pathogens and endogenous ligands, and the requirement of proper UPR for ensuring folding of immunity-related proteins such as cytokines, make these elements essential for sepsis pathogenesis [16], and as such they may serve as sepsis biomarkers.

Our goal was to make use of daily sampling of *tlr4* and *grp78* mRNA levels in septic human patients in order to determine whether their temporal coincidence is a necessary condition for an appropriate response to pathogen. Using clinical and experimental data, we present a mathematical model that reflects the interdependence between the transcription levels oftlr4 and *grp78*. We relate the asynchrony obtained from fitting the data to a damped harmonic oscillation to the patient outcome. Given the importance of GRP78 in modulating the initial response of TLR4 through the control of cytokine production, variations in their expression may be used as an early biomarker for the evaluation of homeostasis maintenance and sepsis progression.

## Results and Discussion

In this study we collected samples of peripheral blood every 24 hours (unless noted otherwise) of sepsis adult patients. An inclusion criterion was that first sample (day 0) was collected as soon as the patient was admitted at the Intensive Care Unit (ICU), and before treatment was initiated. Such inclusion criteria narrowed our cohort considerably, but warranted that gene expression observed in first sample was not disturbed by administered drugs and allowed for an original approach to human sepsis studies. For details on patient recruitment and inclusion/exclusion criteria, refer to Material and Methods section. Samples were collected for up to 5 days, regardless of the clinical outcome of the patient. Clinical and laboratory findings at day of ICU admittance, outcome and respective *causa mortis* are presented in Table 1. Follow-up data was collected at discharge date, at 28 days after ICU admittance, and one year after sepsis event.

**Table 1.** Clinical and laboratory data of sepsis patients

| Patient # | 1 | 2 | 3 | 4 | 5 | 6 | 7 | 8 | 9 | 10 |
| --- | --- | --- | --- | --- | --- | --- | --- | --- | --- | --- |
| Gender | F | M | F | F | M | F | M | M | M | F |
| Age | 32 | 79 | 32 | 89 | 48 | 80 | 43 | 19 | 26 | 81 |
| Infection focus | uterus post-abortion | broncho pneumonia | small bowel colitis, peritonitis | colecistitis | peritonitis | hip prothetic infection, broncho pneumonia | kidney | pericardium and skin abscesses | pleural cavity, pneumonia | septicemia, pneumonia |
| Bacterial culture | N | *C. albicans* + *S. maltophilia* | *S. pyogenes* + *E. coli* | *E. coli* | *E. coli* + fecal Enterococcus | *E. coli* | *S. pyogenes* | *S. aureus* | ND | *S. aureus* + *S. hominis* |
| Comorbidities | N | N | N | Hypertension, previous strokes | N | N | Appendectomy, one month before sepsis | N | N | N |
| C-reactive protein - ICU admission | 179 | 123 | 251 | 172 | 499 | 238 | 61 | 49 | 221 | 181 |
| Leucocytes - ICU admission | 23,400 | 13,800 | 9,800 | 3,900 | 2,600 | 20,200 | 11,200 | 9,500 | ND | 11,700 |
| Neutrophils - ICU admission | 0 | 0 | 1,000 | 1,000 | 1,000 | 6,000 | 5,000 | 1,000 | ND | ND |
| Lactate - ICU admission | 15.9 | 48.1 | 8.3 | 26.2 | 20.6 | 12.9 | 37.4 | 46.9 | 46.9 | 28.1 |
| *Causa mortis* – autopsy results | NA | sepsis and broncho pneumonia | sepsis shock refractory to hemodynamic support | NA | NA | ND | septic shock, multiple organs failure | NA | NA | septic shock, broncho pneumonia, acute kidney failure |
| Outcome | Survival | Death | Death | Survival | Survival | Death | Death | Survival | Survival | Death |

N = pathology absent or absence of bacterial growth *in vitro*. Values in mg/dL units. BD = below detection. ND = not done. NA = not applicable. Normal ranges: C-Reactive Protein (CRP) < 1 mg/dL; leucocytes: 4,000 - 10,000 cells/mm<sup>3</sup>; neutrophils: 1,800 - 7,500 cells/mm<sup>3</sup>; lactate < 20 mg/dL.

Biomarkers used to diagnose sepsis may allow early intervention and consequently may reduce the risk of death. Among these, lactate is largely used, along with a host of other biomarkers that complement its use for early sepsis detection, including C-reactive protein (CRP) and procalcitonin that are produced in response to inflammation [13].

We have determined the CRP and lactate levels for our cohort (Figure 1), in order to predict the patients’ outcome from the onset of sepsis.

**Figure 1.**
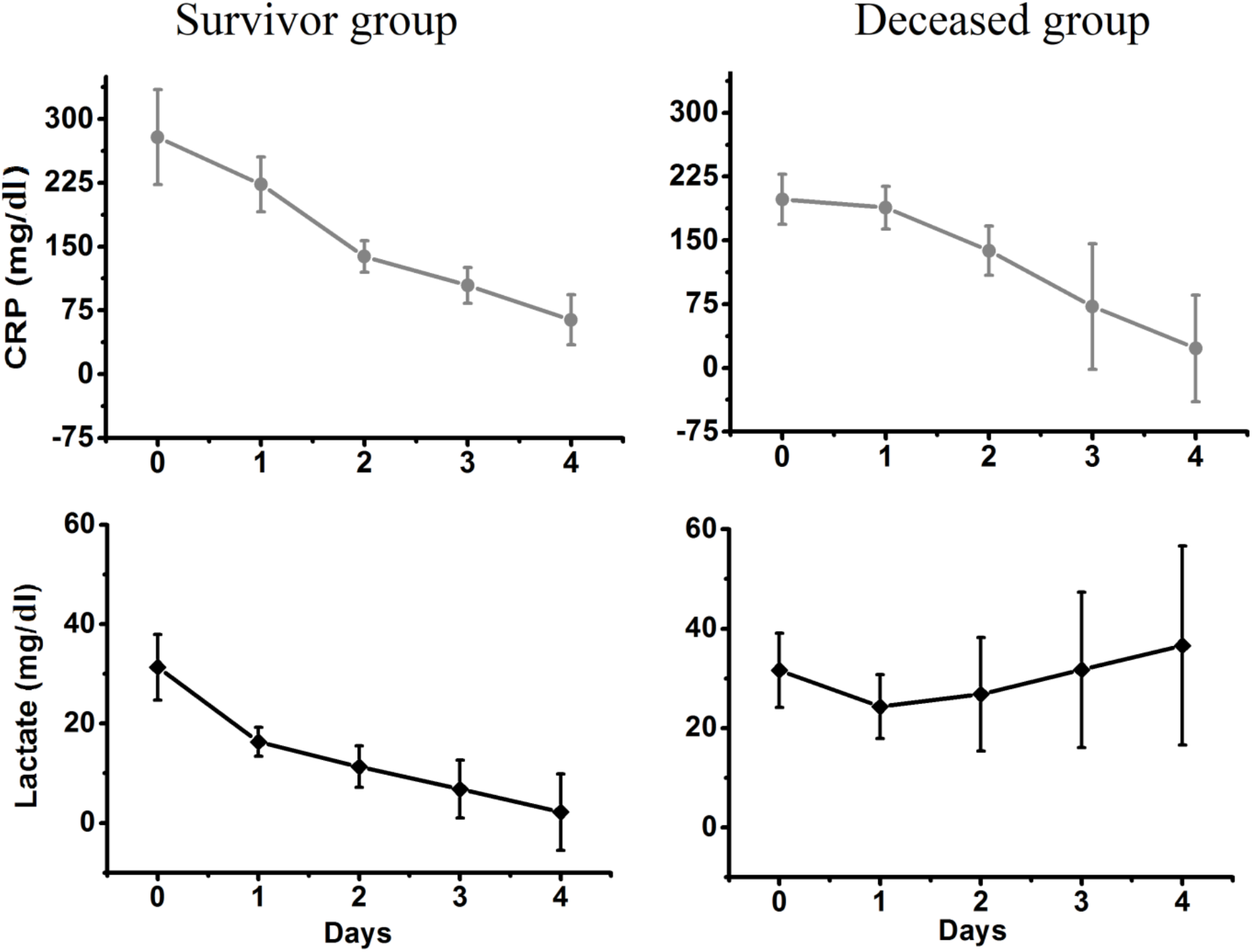
Averaged CRP and lactate time-courses during sepsis progression. Experimental data with standard deviations shown.

Previous studies have revealed average levels of plasma CRP for nonseptic (SIRS) and septic (sepsis, severe sepsis, or septic shock) patients at 7.99 and 11.56 mg/dL, respectively [17]. In our initially non-medicated cohort, average CRP values were 137.8 ± 67.1 mg/dL (survivor group) and 173 ± 88.9 mg/dL (deceased group). For lactate, the accepted threshold of 20 mg/dL [18] is indicative of septic shock is consistent with our mean cohort values of 13.01 ± 8.3 mg/dL (survivor group) and 30.2 ± 4.7 mg/dL (deceased group). While our averaged values at time of ICU admittance are above the clinical limits required to warrant initiation of reactive clinical intervention, they do not in themselves predict the clinical outcome of any patient from our cohort throughout their ICU stay.

In order to test whether gene expression profiles of other molecules involved in sepsis pathogenesis could have prognostic value early upon hospital admission, we focused on *tlr4* and *grp78*. Although whole peripheral blood provides a mixed leukocyte population, which is insufficiently characterized and may have varied significantly among patients and across time, we opted for such samples since (i) their collection and processing reflect real working conditions of intensive care providers at an ICU setting without a cell separation facility, and (ii) every sample’s RNA content was normalized against its own housekeeping gene expression, excluding artifacts of significant variation in RNA content intra- and inter-individuals. Real-time qPCR results are presented in Table 2 as relative quantification units.

**Table 2.** qPCR data from sepsis patients

| Patient # | 1 | 2 | 3 | 4 | 5 | 6 | 7 | 8 | 9 | 10 |
| --- | --- | --- | --- | --- | --- | --- | --- | --- | --- | --- |
| Day 0<br>TLR4<br>(GRP78) | 1.0<br>(1.0)<br>[10 AM] | 1.0<br>(1.0)<br>[11.25 AM] | 1.0<br>(1.0)<br>[11 AM] | 1.0<br>(1.0)<br>[11.30 AM] | 1.0<br>(1.0)<br>[10 AM] | 1.0<br>(1.0)<br>[4 PM] | 1.0<br>(1.0)<br>[10 AM] | 1.0<br>(1.0)<br>[12 AM] | 1.0<br>(1.0)<br>[4 AM] | 1.0<br>(1.0)<br>[11 PM] |
| Day 0.5<br>TLR4<br>(GRP78) |  |  |  | 0.6<br>(0.64)<br>[11.30 PM] |  | 1.97<br>(1.7)<br>[4 AM] | 4.3<br>(0.66)<br>[10 PM] | 0.56<br>(3.6)<br>[12 PM] | 1.97<br>(0.57)<br>[4 PM] |  |
| Day 1<br>TLR4<br>(GRP78) | 0.3<br>(1.23)<br>[10 AM] | 0.2<br>(0.14)<br>[11.30 AM] | 2.5<br>(1.7)<br>[11 AM] | 0.13<br>(1.5)<br>[11.30 AM] | 0.89<br>(1.05)<br>[10 AM] | 1.7<br>(1.6)<br>[4 AM] |  | 1.4<br>(3.07)<br>[12 AM] | 1.71<br>(0.5)<br>[4 PM] | 1.63<br>(1.18)<br>[11 AM] |
| Day 2<br>TLR4<br>(GRP78) | 0.4<br>(2.0)<br>[10 AM] | 1.3<br>(0.74)<br>[11.30 PM] | 1.5<br>(1.4)<br>[11 AM] | 0.11<br>(0.9)<br>[11.30 AM] |  |  |  | 1.01<br>(2.8)<br>[10 AM] | 1.0<br>(0.44)<br>[4 PM] | 1.43<br>(0.8)<br>[1 PM] |
| Day 3<br>TLR4<br>(GRP78) | 0.7<br>(1.24)<br>[2 PM] |  | 0.63<br>(1.4)<br>[11 AM] | 0.25<br>(0.28)<br>[11 AM] |  | 0.98<br>(3.4)<br>[4 AM] |  | 1.7<br>(4.0)<br>[10 AM] | 0.45<br>(0.45)<br>[4 PM] |  |
| Day 4<br>TLR4<br>(GRP78) | 1.0<br>(0.88)<br>[10 AM] |  | 0.9<br>(16.4)<br>[11 AM] |  |  | 0.45<br>(0.4)<br>[4 AM] |  | 1.6<br>(4.0)<br>[10 AM] | 0.44<br>(0.64)<br>[4 PM] |  |
| Day 5<br>TLR4 |  |  |  |  |  | 0.44<br>(0.51) |  |  |  |  |
| (GRP78) |  |  |  |  |  | [4 AM] |  |  |  |  |
| Outcome | Survival | Death | Death | Survival | Survival | Death | Death | Survival | Survival | Death |

Day 0 refers to admission to ICU and diagnosis of sepsis. Subsequent sampling was performed every 12 (“day 0.5”) or 24 hours. Times of sampling are indicated in brackets. qPCR values expressed as relative quantification units, *tlr4* in the 1<sup>st</sup> line, *grp78* within round parentheses in the 2<sup>nd</sup> line.

Although conventional statistical analysis could detect significant differences between values from survivors and deceased patients (p-values of 0.01 (*tlr4*) and 0.02 (*grp78*), refer to Materials & Methods section for details), it does not yield information on the dynamical changes of these two parameters, neither has predictive value for the ICU clinician. To this end, we have studied the interrelation between the transcriptional profiles of *grp78* and *tlr4* using a damped harmonic oscillation model. The equation describing an energy dissipating oscillator is used to describe a system that is able to accommodate exogenous (environmental) and endogenous (*i.e*. fluctuations in mRNA or protein levels) inputs without expanding its output limits. Such a system is able to resist to extra- and intra-cellular perturbations without changing the nature or intensity of its output [19]. NFκB is one example of a damped oscillator, where given a constant source of TNF-α(input), the amplitude of the NFκB response decreases gradually until cessation [20]. Gene transcription in a human osteosarcoma cell line was also shown to behave as a damped oscillator, where transcription occurred in bursts with decreasing amplitude and frequency [21]. In sepsis pathogenesis, this could be illustrated by the patient who is able to keep cytokine synthesis under control, preventing the deleterious effects of the “cytokine storm” that corresponds to an increase of its response. Regardless of the endotoxin concentration (exogenous input), the system is able to respond within the limits of a homeostatic inflammatory response, and return to the initial conditions of homeostasis. A dissipating system is thus able to “release” the excessive input while restraining the output within energetically-optimized limits. One of the mechanisms to achieve this fine-tuning is by oscillating the expression of repressor or inducer elements in the network, adjusting the intensity of the response in each cycle [21].

Averaged (main panels) and raw (inserts) daily profiles of *tlr4* and *grp78* transcript levels of patients are presented in Figure 2.

**Figure 2.**
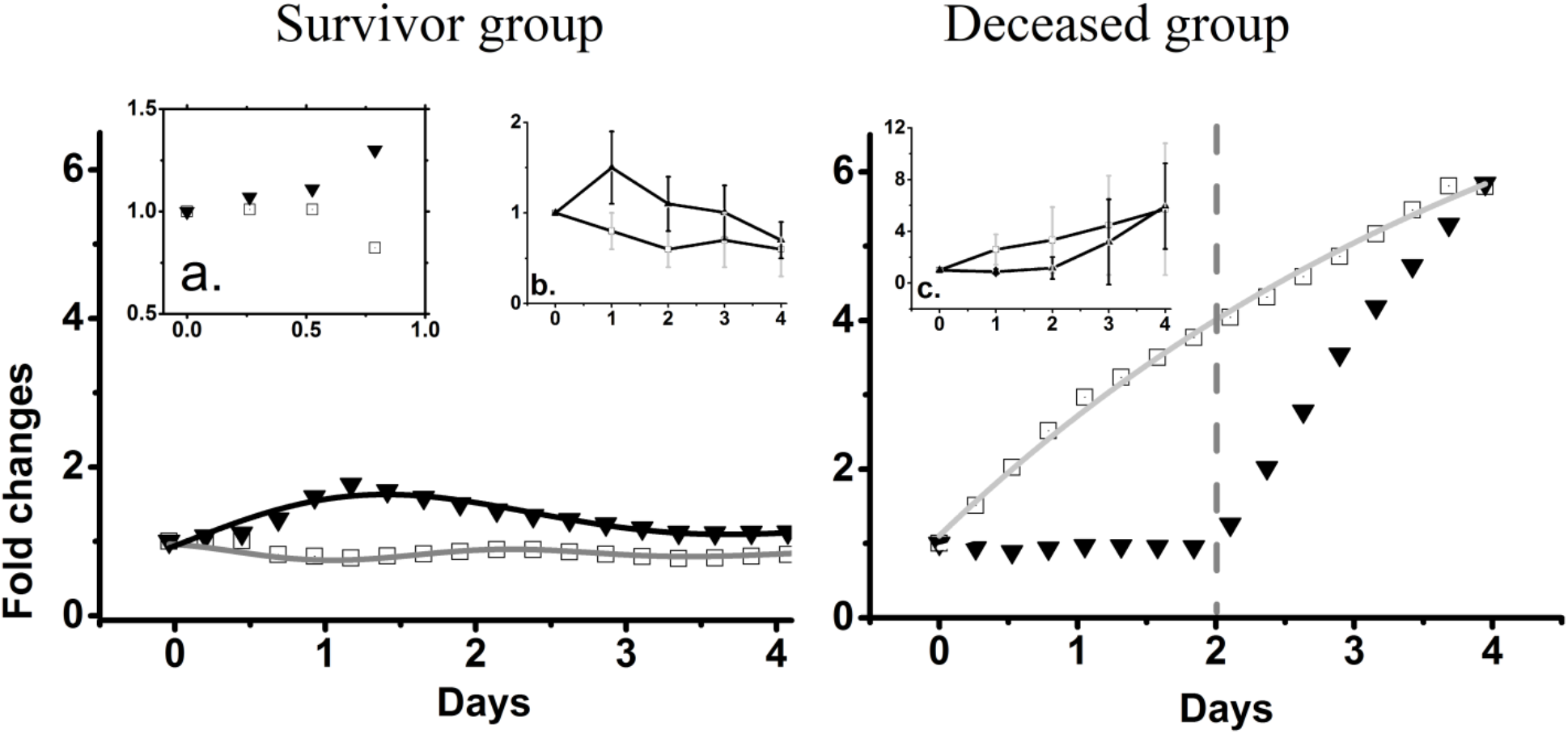
Averaged mRNA time-courses during sepsis progression. Experimental data and fitted curves for the survivor group (left panel) and deceased group (right panel) Squares represent *tlr4*, triangles represent *grp78*. Fitted curves for transcripts of the two genes are depicted as a line. For *grp78* transcripts in deceased patients (triangles, right panel), we have considered only the sinusoid part of the curve until the switch point at approximately 48 hours (refer to text for further details). The insert (a.) shows the first 24 hours of averaged data in the survivor group. Inserts (b.) and (c.) show the raw data with standard deviations bars (no interpolation).

In patients who survived sepsis (Fig 2, left panel), the accumulation of transcripts for both *tlr4* and *grp78* presents an initial synchronized increase during the initial 6 hours (see figure insert (a.) for details). This is followed by a marked down-regulation of *tlr4* (square symbols) until 48 hours, and by an increase of *grp78* (triangle symbols) that peaks around 24 hours, with subsequent continuous decrease. In deceased patients (Fig 2, right panel), *tlr4* values (square symbols) increase exponentially, while *grp78* (triangle symbols) mRNA from deceased patients oscillate until an exponential increase at a switch point at day 2.

Fluctuations in the transcript levels of *tlr4* (survivor group only) and *grp78* (both groups) were simulated using an equation describing an energy dissipating harmonic oscillator:

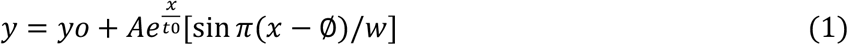

Data were fitted to a damped sine wave by varying the following parameters: y_o_ = offset, *A* =initial amplitude, t_0_ = decay constant, ø = phase shift and w = period, as shown in Table 3.

**Table 3.**
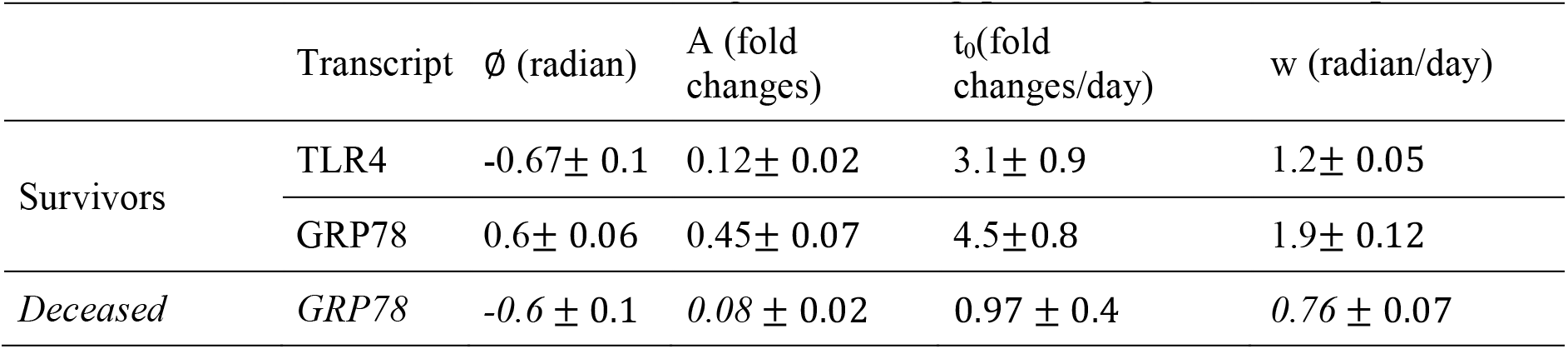
Fitted values to the curves describing the tlr4and *grp78* changes in transcripts

| | Transcript | $\phi$ (radian) | A (fold changes) | $t_0$(fold changes/day) | w (radian/day) |
| --- | --- | --- | --- | --- | --- |
| Survivors | TLR4 | $-0.67 \pm 0.1$ | $0.12 \pm 0.02$ | $3.1 \pm 0.9$ | $1.2 \pm 0.05$ |
| | GRP78 | $0.6 \pm 0.06$ | $0.45 \pm 0.07$ | $4.5 \pm 0.8$ | $1.9 \pm 0.12$ |
| Deceased | GRP78 | $-0.6 \pm 0.1$ | $0.08 \pm 0.02$ | $0.97 \pm 0.4$ | $0.76 \pm 0.07$ |

The average *tlr4* mRNA values of deceased patients increase exponentially from day 0and thus, were fitted to the following asymptotic exponential equation:

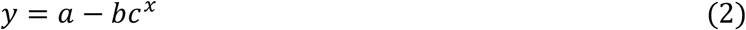

with fitting parameters a = asymptote (9.2 ± 0.8), b = response range (8.1±0.7), and c = rate of change (0.84 ±0.03).

Previous measurements of *tlr4* mRNA production in murine macrophages yielded a high peak around 20 hours post-stimulation, followed by a smaller peak around the 40 hours mark [22]. While the first peak is absent in the *tlr4* transcript variations of our survivor patients group, the second peak is recovered, as evidenced through the w-parameter. The increase of *trl4* transcripts on the second day upon ICU admission in survivors may signal the presence of a robust regulatory mechanism that limits the initial amount of available TLR4 so as to dampen the subsequent inflammation stage. In contrast, in the deceased group, loss of regulation appears to lead to an exponential increase in the production of *tlr4* mRNA, expanding the limits of the inflammatory response with lethal effects for the patients. As endotoxin concentration increases the initial production and the subsequent half-life of *tlr4* mRNA [23], we hypothesize that a more robust and physiological response to the septic challenge may also require a rapid regulation to the subsequent increase in TLR4 expression and activity, chiefly through controlled transcription of *tlr4* and increased transcription of its different modulators, including GRP78.

We observed the opposite trend for the transcripts of *grp78*, reflecting its role as an importantTLR4 regulator: in deceased patients, within the first 48 hours (to which data could be fitted), the amplitude and decay constants are lower than those found in the survival group, reflecting lower levels of *grp78* expression in deceased patients when compared to patients that survived. The exponential increase in *grp78* transcripts past this point in the deceased group may relate to the findings of higher levels of extracellular isoforms of the heat shock protein-70kDa (Hsp70) family, including GRP78 [24], that are implicated in the progression to septic shock, in the subsequent immunosuppression phase of sepsis [25], and are found in the serum of septic patients [26]. It is possible that the oxidative stress caused by engagement of TLR4 and cytokine receptors may contribute to the modulation of serum Hsp70 levels, by causing its active secretion [27], or release through necrosis [28]. Furthermore, cell-membrane localized GRP78 assist in recruiting oxidized phospholipids that promote both endothelial barrier integrity against refractory hypotension [29] or rescue from TLR4-based acute inflammation [30]. While extracellular Hsp70 act as an agonist to TLR4, intracellular members of Hsp70 family markedly inhibit not only signaling downstream of TLR4, but also modulate its ubiquitination and proteasomal degradation [31].

Recognition of pathogens and subsequent activation of inflammatory signaling was proposed to be regulated by a “macrophage-intrinsic clock”, affecting TLR4 circadian activity regulation through in-phase gene transcription of downstream modulators radio-protective 105 (RP105) and myeloid differentiation factor-1 (MD1), along with GRP78 itself [32]. Within the survivor group, we measured an inverse relation between the higher amplitude of *grp78* transcripts accumulation with respect to *tlr4*, and comparatively similar decay rates. Furthermore, the phase delay for *tlr4* transcription succeeded the corresponding transcription of *grp78* in survivors, with the inverse trend observed in the group of deceased patients. Other important TLR4 modulators, including MD2 and Myeloid differentiation primary response gene 88 (MyD88), are also upregulated upon LPS stimulation of human monocytes and macrophages [23]. Moreover, dysregulation of central and peripheral circadian rhythms was noted in studies of septic patients in ICU, and complete loss thereof was characteristic of non-survivors [33]. A damped oscillator system predicts the existence of a frequency modulator that acts as a common regulatory mechanism [34]. One candidate as frequency modulator of the mutual relationship between the expression of *tlr4* and *grp78* is interleukin-10 (IL-10). IL-10 signaling in unstressed cells blocks *grp78* transcription by inhibiting nuclear translocation of transcription factor ATF-6 and its binding to *grp78* promoter [29]. IL-10 upregulates the expression of TLR4, MyD88 and CD14 [36]. Upon LPS treatment, *il10* mRNA expression has a very slow onset and is not detected in culture before a 6 hours mark, increasing steadily to 24 hours after stimulation [37]. Both IL-10 mRNA and protein responses are slow to occur in human monocytes [38] and microglial cells [39]. It is possible that this delayed response may contribute to the initial *grp78* upregulation that we observed in the survivors group. As IL-10 progressively accumulates up to 48 hours [37], the *tlr4* expression may become dominant. However, increasing IL-10 levels upon TLR4 signaling also inhibits the induction of the microRNA miR-155, that results in higher concentrations of Src homology 2 (SH2) domain-containing inositol-5′-phosphatase 1 (SHIP1), culminating with further inhibition of TLR4 own signaling [40]. IL-10 is an anti-inflammatory modulator responsible for the dampening of the initial pro-inflammatory response through autocrine/paracrine feedback mechanisms. As such, TLR4-mediated induction of IL-10 may be subject to intrinsic temporal regulation that delays IL-10 production to allow time for the initiation of the immune response upon LPS stimulation [41]. The combination of positive and negative regulatory loops involving IL-10 may be responsible for the oscillatory expression patterns we measured in our cohort.

Constructing gene expression profiles for key modulators of the sepsis syndrome represents a novel and non-invasive means to assess its evolution. Such gene expression profiles of mediators have been previously introduced for other pathological conditions, such as coronary artery disease [42] or, using TLR7 and TLR8, for acute ischemic stroke [43]. Taken together, our observations along with retrospective studies on the times of sepsis deaths [14] may reflect the requirement for TLR4 modulators such as GRP78 to be timely expressed before TLR4 upregulation, in order to prevent subsequent excessive pro-inflammatory signaling.

Although our cohort is small, effect size calculation showed large effects (1.868 for *grp78* and 1.208 for *tlr4*). Our findings suggest that sepsis prognosis may be inferred early upon admission if on the first day of hospitalization three blood samples are collected so as to establish the amplitude, decay constants and phase shifts of *tlr4* and *grp78* mRNA accumulation in patients’ peripheral whole blood. In our dataset, higher levels of *grp78* and lower levels of tlr4mRNA anticipated a positive evolution of sepsis, leading to patient survival. In a follow-up one year after samples were collected, none of the survivors presented any morbidity related to sepsis. In contrast, intense accumulation oftlr4 transcripts accompanied by very low (if any) levels of *grp78* mRNA in the first two days after sepsis onset reflected a poor prognosis. All deceased patients died within a period of 28 days upon ICU admission while all survivors were still alive at a follow-up after one year, confirming predictions made by our model using data from the first 5 days of hospitalization. It is an intriguing idea that some crucial genes should be kept under tight transcriptional control if the organism is to maintain homeostasis and survive. Besides being a useful marker of homeostasis breakage the loss of transcriptional control might, *per se*, be a driving force behind the pathological state in sepsis. If confirmed by others, new venues of research shall open, aiming at understanding and preserving transcriptional control during intense inflammatory conditions such as SIRS and sepsis. We envision that model refining through multicentric collection of additional *ex vivo* data will shed more light on the mechanisms regulating gene transcription during intense systemic inflammatory conditions, and will provide the intensive care personnel an additional but very sensitive compass to guide subsequent therapy.

## Material and Methods

### Patient recruitment

The study involved consecutive data collection from clinical and surgical patients, aged over 18 years old, referred to the ICU of the University Hospital at the University of São Paulo, and diagnosed with sepsis according to Acute Physiology and Chronic Health Evaluation II (APACHE) and SOFA within the first 24 hours of admission. During the ICU stay, all patients received medical treatment according to accepted guidelines [44].

Peripheral blood samples as well as clinical and laboratorial data were collected at patient admission to the ICU within the first 24 hours of sepsis onset and every 24 hours up to five days. The total length of the hospital stay and outcome were recorded. Treatment with corticosteroids and/or antibiotics in the days previous to admission to the ICU was an exclusion criterion for recruitment. Additional information was also collected, including comorbidities, smoking and alcoholism history. The Institutional Research Ethics Committees (CEP-HU/USP 940/09 SISNEP CAAE 0048.0.019.198-09, CEPSH-ICB 975/10) approved of the study and patients signed a written informed consent prior to enrollment in the study.

### Laboratorial investigations

Sites of suspected or documented infection, blood gas analysis, complete blood count, C-reactive protein, and microbiological cultures were documented (Table 1). Subsequent daily measurements of all these parameters were performed and may be provided on request. Serum levels of total bilirubin and fractions, creatinine, urea, sodium, potassium and glucose were also measured (data not shown).

### Transcripts quantification

Two and a half ml of peripheral blood were collected into PAXgene RNA tubes (PreAnalytiX BD) every 24 hours from time of admission (unless stated otherwise), up to five days (or less, in those cases where the patient died or was discharged). Tubes were kept at -20°C for up to 4 months until RNA isolation. Total RNA was isolated using PAXgene Blood RNA kits (Qiagen, São Paulo, Brazil), and synthesis of cDNA was performed in the presence of RNase inhibitor using random primers and the High Capacity cDNA Reverse Transcription kit (Thermo Fisher Scientific, São Paulo, Brazil), according to the manufacturer’s instructions. Real-time quantitative PCR (qPCR) were performed using specific primers and Fast SYBR^®^ Green Master Mix (Thermo Fisher Scientific, São Paulo, Brazil) in an Agilent Mx3005P qPCR System (Agilent Technologies, Santa Clara, CA, USA). Triplicate reactions were run for each sample. Oligonucleotides used for transcripts quantification: GRP78/BiP forward (NM_005347.4, exon 1) 5′- CGAGGAGGAGGACAAGAAGG, GRP78/BiP reverse (NM_005347.4, exon 3) 5′- CACCTTGAACGGCAAGAACT, GAPDH forward (NM_001289746.1, exon 7) 5′- CGACCACTTTGTCAAGCTCA, GAPDH reverse (NM_001289746.1, exon 8) 5′- CTGTGAGGAGGGGAGATTCA, TLR4 forward (NM_138557.2, exon 1) 5′- CGCTTTCACTTCCTCTCACC, TLR4 reverse (NM_138557.2, exon 1-2) 5′- ATTAGGAACCACCTCCACGC. Cycling conditions were: 95°C for 20 secs, followed by 40 cycles of 95°C for 3 seconds and 60°C for 30 seconds. Melting curves were acquired at 95°C. Reaction efficiencies ranged between 95 and 97%. Transcripts levels were normalized to *gapdh* mRNA and the mean relative abundance values were calculated using the 2-ΔΔCt method [45], and are shown in Table 2.

### Data analysis

*tlr4* and *grp78* mRNA data was averaged with Origin 9.1 (OriginLab, Northampton, MA, USA), using a linear interpolation algorithm. The resulting averaged curves were fitted to a damped sine wave equation by manually varying the fitting parameters until 0.9 <R^2^< 1. Data for *grp78* mRNA from patients who succumbed were fitted to the first 48 hours only.

### Statistical analyses

Data was tested for normality at 5% decision level using the Kolmogorov-Smirnov normality test. Comparisons among treatment groups were performed using Fisher’s exact test for pair-wise correlations between the averaged values of the survivor and the deceased patient groups, within either category of gene transcripts, yielding p-values of 0.01 (*tlr4*) and 0.02 (*grp78*). Between the deceased and the survivors, power was 0.70 (*tlr4*) and 0.67 (*grp78*), at 95% confidence level, with α = 0.05. Effect size was calculated using Cohen’s d equation, survivors vs. deceased, yielding values of 1.868 (*tlr4*) and 1.208(grp78). All calculations were performed in Origin 9.1 (OriginLab, Northampton, MA, USA).

## Acknowledgements

We thank Drs. Júlio Aliberti, Lisl Shoda, Luciana Moraes, Giorgio Trinchieri and Grégoire Altan- Bonnet for the critical reading of this manuscript.

